# Improved Wound Closure Rates and Mechanical Properties Resembling Native Skin in Murine Diabetic Wounds Treated with a Tropoelastin and Collagen Wound Healing Device

**DOI:** 10.1101/2020.10.01.322636

**Authors:** Robert S. Kellar, Robert B. Diller, Aaron J. Tabor, Dominic D. Dominguez, Robert G. Audet, Tatum A. Bardsley, Alyssa J. Talbert, Nathan Cruz, Alison Ingraldi, Burt D. Ensley

## Abstract

Chronic wounds in patients suffering from type II diabetes mellitus (DMII) where wounds remain open with a complicated pathophysiology, healing, and recovery process is a public health concern. Normal wound healing plays a critical role in wound closure, restoration of mechanical properties, and the biochemical characteristics of the remodeled tissue. Biological scaffolds provide a tissue substitute to help facilitate wound healing by mimicking the extracellular matrix (ECM) of the dermis. In the current study an electrospun biomimetic scaffold, wound healing device (WHD), containing tropoelastin (TE) and collagen was synthesized to mimic the biochemical and mechanical characteristics of healthy human skin. The WHD was compared to a commercially available porcine small intestinal submucosa (SIS) matrix that has been used in both partial and full-thickness wounds, Oasis^®^ Wound Matrix. Wound closure rates, histochemistry, qPCR, and mechanical testing of treated wound sites were evaluated. The WHD in a splinted, full-thickness, diabetic murine wound healing model demonstrated an enhanced rate of wound closure, decreased tissue inflammation, skin organ regeneration, and a stronger and more durable remodeled tissue that more closely mimics native unwounded skin compared to the control device.

## Introduction

Chronic, non-healing, or slow to heal wounds present a significant and growing health problem in the United States, with an estimated 6.5 million people affected, at an annual cost of US $20 billion, with the highest risk groups represented by the elderly and the increasing prevalence of lifestyle diseases such as diabetes and obesity (Frykberg 2015, Sen, 2019). These wounds can take years or decades to heal causing significant emotional distress and physical pain to the patient, including their families (Jarbrink 2017). Chronic diabetic wounds pose pathophysiological abnormalities compared to “normal” wound healing (Falanga, 2005).

The current numbers reported by the Center for Disease Control (CDC) estimates that more than 122 million people in the United States are living with diabetes (34.2 million) or prediabetes (88 million) (CDC, 2020). Of the more than 23 million people in the US affected by diabetes, 10-15% will have at least one diabetic foot ulcer (DFU) in their lifetime (Margolis, 2011), resulting in more than $176 billion per year in health care expenditures (Mathieu, 2006 and Menke, 2007).

The current standard of care for diabetic wounds varies depending on the site of the wound, type of infection and clinician applying the treatment. Many treatment interventions include the application of wound dressings, surgical procedures, pressure relief therapy, biologics, antibiotic regimens, hyperbaric chambers, negative pressure devices and regenerative medical therapies. Wound dressings provide and maintain an optimal humid environment for the wound bed and subsequent healing (Hingorani, 2016). Surgical interventions include debridement, ridding the wound bed and surrounding wound margins of necrotic tissue (Chharbra, 2017), (Hingorani, 2016), and pressure relief applications which use casting and individualized shoe inserts or post-operative shoes to reduce mechanical stress to the wound (Frykberg, 2006), (Health Quality Ontario, 2017). Biologic therapies include antimicrobial peptide applications (Santos 2016), (Gawande, 2014), (Saeed, 2013), (Lipsky, 2008) and a revival of larval treatment, which debride the wound without surgical involvement (Lipsky, 2008), (Sherman, 2003), (Hassan, 2014). Antibiotic regimens include a wide array of mixtures and classes, such as, glycopeptides and beta-lactams, although the infectious agent generally establishes the antibiotic regimen (Bergman, 2016), (Singh, 2017). Hyperbaric chambers provide a rich supply of oxygen to hypoxic wounds and have been demonstrated to promote fibroblast proliferation, stimulate angiogenesis and increase the immune response, which are all requisite for progression of the wound healing cascade (Han, 2017). Negative-pressure devices, also called wound-vacs, have produced positive impacts on wound healing. These vacuum assisted devices, keep the wound bed moist, remove exudate and bring a fresh supply of blood to the wound bed (Han, 2017). Lastly, tissue engineering is a field within regenerative medicine aimed at restoring the native properties of the injured tissue. Within this field, biomimetic scaffolds have been fabricated, or preexisting tissue has been manipulated to create a device that mimics the extracellular matrix (ECM). These biomimetic ECM devices can then be implanted into the wound bed aiding in the restoration of healthy tissue. Commercially available examples of these ECM fillers include Biobrane, Integra, AlloDerm and a porcine intestinal sub-mucosa membrane titled Oasis^®^ Wound Matrix (Oasis SIS Wound Dressing II (K993948)). However, many of these products, along with other commercial wound devices, rely on harvested animal or cadaver tissue for production. Processes used in tissue harvesting for many of these commercial devices, including decellularization, subject the tissue to invasive processing which alters and disrupts the native ECM proteins and structure, potentially impacting their ability to promote progression of the healing cascade in a chronic wound (Badylak, 2015).

When an injurious event occurs, normal wound healing follows an overlapping, three-phased approach (Clark, 1996). Phase one includes inflammation, followed by the proliferative/migration phase and lastly the tissue remodeling phase. During a diabetic wound healing response, the three-phases occur but with pathophysiologic changes. During the inflammatory phase patients with DMII demonstrate a disequilibrium of essential cytokines such as interleukins (ILs), tumor necrosis factor-alpha (TNF-α), PDGF, epidermal growth factor (EGF) among others (Chhabra, 2017) (Werner, 2003) (Xioa, 2014), (Pradhan, 2009). Additionally, the leukocyte exocytotic cytokine release is altered increasing the chance for infection (Xioa, 2014), (Pradhan, 2009). During the proliferative/migratory phase, angiogenesis brings in a fresh supply of blood and oxygen to the wound bed. Next, fibroblasts proliferate and aid in the formation of the ECM, which contains collagens, fibronectin, vitronectin and elastin. This ECM allows for the migration of additional cells, such as neutrophils, macrophages, keratinocytes, dendritic cells, to the injured area allowing for reepithelization and remodeling to occur (Singer & Clark, 1999). In a diabetic wound, the elevated hyperglycemic levels alter the migratory capacity of these proliferating cells, causing a deficiency in the new ECM deposition and recruitment of epithelial cells (Santoro, 2005), (Lan, 2008), (Maione, 2016). These cumulative events lead to a chronic or slow-to-heal diabetic wound.

The quantity and presence of collagen I and elastin, play an important role in the proliferative phase of wound healing. In the early stages of a wound, damage to the dermis signals recruitment of macrophages which secrete MMP-12 (elastase) that generates elastokines from resident elastin. These small elastin peptides signal a transient increase in inflammation associated with neutrophil infiltration and migration signals, furthering progression through the inflammatory and remodeling phases in wound healing in healthy tissue (Almine, 2013). Supplying the wound bed with physiologically relevant amounts and ratios of collagen I and elastin facilitates progression of the healing cascade. Therefore, we have developed bioengineered scaffolds using human skin proteins in an architectural orientation that mimics the ECM, demonstrating improved cutaneous wound closure *in vivo* alone, or when incorporating therapeutics such as platelet-rich plasma, stem cells, and growth factors that promote accelerated wound healing compared to control therapy (Machula, 2014), (Tabor, 2016). In this study we hypothesized that a wound healing device (WHD) comprised of human collagen I and tropoelastin in physiologically relevant proportions would encourage a chronic wound halted in the inflammatory phase to progress into the remodeling phase. To test this hypothesis, full thickness, splinted wounds in diabetic mice were treated with biomimetic WHDs to assess wound healing and the resulting remodeled tissue (scar) mechanical properties.

In this study, the WHD demonstrated an enhanced rate of wound closure, decreased tissue inflammation, skin organ regeneration, and a stronger and more durable remodeled tissue that more closely mimics native unwounded skin compared to the control devices. The WHD has the potential to provide a novel, commercially available device for the treatment of chronic non-healing diabetic wounds.

## Materials and Methods

### Wound Healing Device (WHD) creation

Scaffolds were created using human dermal collagen type I (VitroCol, Advanced Biomatrix, Carlsbad, CA) and recombinant human dermal tropoelastin (rhTE, Protein Genomics, Sedona, AZ). The proteins were dissolved into 1,1,1,3,3,3-hexafluoro-2-propanol (HFIP, Oakwood Chemical, Estill, SC). The biomimetic WHDs were prepared using a ratio of 9:1 collagen to rhTE. The solution was loaded into a 5 mL syringe and dispensed with a syringe pump at a constant rate of 3mL/hr. A voltage of 24-27kV was used to create circular scaffolds onto a circular aluminum foil target. The scaffolds were dried in a desiccator at room temperature overnight. The scaffolds were then sterilized by exposure to ultraviolet (UV, 254 nm) light for one hour on each side and prepared using a sterile 8mm biopsy punch.

### *In vivo* diabetic wound healing assay using a splinted wound model

Murine strain (C57BKS.Cg-m+/+*Lepr^db^/*J mice (db/db), Jackson Labs, Bar Harbor, ME) animals were used in all the experiments. All animals were handled in accordance with the Institutional Animal Care and Use Committee of Northern Arizona University, and the National Research Council Guide for the Care and Use of Laboratory Animals (National Research Council, 2011)). The murine diabetic model was used following Northern Arizona University (NAU) IACUC approval, Protocol #12-006R1. Seventy-two (72) 6-8-week-old mice, 36 male and 36 female, were placed into two groups. See Table 1 for experimental layout. The study followed a 14-day and 28-day research plan: day 0 = full thickness wound creation; day 14 and day 28 = euthanasia. All animals survived the duration of the evaluation.

**Table 1.** Description of the grouping and animal use for evaluation.

| Group | Timepoint | Number of Animals (2 wound sites per animal) |  |  |  |  |  |
| --- | --- | --- | --- | --- | --- | --- | --- |
|  |  | Mechanical testing |  |  | Biological testing (Histology/IHC/qPCR) |  |  |
|  |  | Treatments |  |  | Treatments |  |  |
|  |  | Control | Oasis® | WHD | Control | Oasis® | WHD |
| 1 | 14 Days | 3♀ 3♂ | 3♀ 3♂ | 3♀ 3♂ | 3♀ 3♂ | 3♀ 3♂ | 3♀ 3♂ |
| 2 | 28 Days | 3♀ 3♂ | 3♀ 3♂ | 3♀ 3♂ | 3♀ 3♂ | 3♀ 3♂ | 3♀ 3♂ |

### Surgical preparation

Mice were anesthetized in a closed chamber using 2-2.5% isoflurane with 0.5-1 L/min oxygen to effect, evidenced by lack of response to toe pinch stimulus. Mice were placed on a towel-covered heating pad to maintain normal core body temperature while under anesthesia until fully recovered. Mice were transferred to a preoperative field and connected to a nose cone delivering 2% isoflurane in 1 L/min oxygen. Mice were weighed and blood glucose levels were measured by collecting a droplet of blood by tail snip and reading in a Contour Next EZ Meter using Contour Next test strips (Ascensia Diabetes Care, Parsippany, NJ). The tail snip was sealed with a droplet of super glue. Mice were then shaved and depilated with Nair™ (Church and Dwight, Ewing, NJ) to obtain a smooth skin surface. Following saline rinse to remove Nair, mice received a subcutaneous pre-operative dose of buprenorphine (0.01mg/kg) and 1 cc sterile saline. Mice were transferred to a sterile surgical field and connected to a nose cone delivering 1.5-2% isoflurane in 1 L/min oxygen to maintain anesthesia.

### Surgical procedure

A randomization scheme was used to enroll animals into treatment and control groups according to Table 1. Wounds were created following previously published methods (Galiano et al., 2004; Machula et al., 2014), briefly described here. The dorsum was scrubbed with betadine solution and two full thickness wounds per dorsum were created lateral to the midline by scoring the skin with a sterile 8 mm biopsy punch and excising the tissue with sterile iris scissors. A sterile, donut-shaped silicone splint (Grace Bio-Labs, Bend, OR) was centered over the wound and secured with six interrupted 6-0 polypropylene sutures. The two treatments; WHD and Oasis^®^ (Smith+Nephew, Fort Worth, TX) were prepared using a sterile 8 mm biopsy punch. The treatment devices were added to the splinted wound and allowed to uniformly contact the wound bed while the control sites remained untreated. All wounds were photographed before and immediately after treatment. All wounds from all treatment groups were covered with Tegaderm^®^ dressing (3M, St. Paul, MN) and the mice were individually caged until fully recovered.

### Postoperative care and wound photography

Mice were housed individually to prevent animal interference with wound sites. Cages were placed on a static ventilation caging system and welfare checks were performed once daily for normal activity, food and water consumption, excretion and grooming. Postoperative analgesics were administered within the first 1-2 days as needed.

Every other day, mice were anesthetized with 2% isoflurane in 1 L/min oxygen in a chamber, then transferred to a prep area remaining under anesthesia. The Tegaderm was carefully removed and visual observations of the wound were recorded, followed by photography of the wound bed. A new piece of Tegaderm was applied and mice were recovered in their cage, then returned to their housing room.

### Wound closure analysis

Wound closure was quantified by measuring the open area of the wound every 2 days through the duration of healing. The images in each study were analyzed blinded to treatment and sex. Percent wound closure was analyzed with NIH ImageJ, v1.52a (Bethesda, MD) by standardizing each image using the millimeter ruler captured in each photo. The open wound margin was traced with a freehand tool, generating an area in mm^2^. The wound bed for each day was normalized to time 0 using the following equation a:

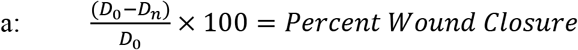

where *D*_*0*_ = Initial wound area measurement and *D*_*n*_ = Day of wound area measurement. The data were then identified by sex and treatment for analysis.

### Tissue collection

After 14 or 28 days, the animals were euthanized with 5% isoflurane, followed by cervical dislocation. The skin of the dorsum was cut on 3 sides to create a skin flap. The wound bed was excised from the underside. The samples for immunohistochemistry (IHC) and histology were cut out using a 10 mm biopsy punch, then sectioned in half, coronally. One half was placed in 4% paraformaldehyde in 1x PBS at 4°C for histology and IHC and the other half was placed in ice cold RNAlater at 4°C for PCR analysis. The samples for mechanical testing were prepared by cutting the samples into dog-bone templates (ASTM D412, Gallagher et al., 2012) and stored in KREBS until tested.

### Histology and Immunohistochemistry

Tissue samples were removed from fixative after 24 hours at 4°C and transferred to cold 70% ethanol until ready for processing. The tissue was dehydrated with graded ethanol to paraffin. The tissue was sectioned at 5 μm and stained with hematoxylin and eosin. These slides were used to evaluate the wound healing response and re-epithelialization.

The tissues were also processed for IHC (Histotox, Boulder CO). All tissues were processed on a Leica Bond Rxm using standard chromogenic methods. For antigen retrieval, slides were incubated in pH 9 EDTA-based buffer for 2 hours at 70°C for CD31 (endothelial cells) and CD163 (monocytes and macrophages) or incubated with Proteinase K for 8 minutes at room temperature for elastin (Lau et al., 2004). Slides were incubated with appropriately diluted antibody for 30 minutes for CD31 (1:500, Abcam ab182981) and CD163 (1:400, Abcam ab182422) or for 45 minutes for elastin (1:200, Abcam ab23748). Antibody binding was detected using an HRP-conjugated secondary polymer, followed by chromogenic visualization with diaminobenzidine (DAB). A Hematoxylin counterstain was used to visualize nuclei.

All slides were digitally scanned using a Hamamatsu NanoZoomer (Hammamtsu, Japan) with a resolution of 0.5 μm per pixel and analyzed using ImageScope software and algorithms (Leica, Buffalo Grove, IL). The inflammatory response was evaluated using a nuclear counting algorithm, while the presence of microvasculature was evaluated using a microvessel analysis algorithm and elastin was evaluated using the color deconvolution algorithm.

### Gene expression

Tissue samples for gene expression analysis were stored at 4°C for 24 hours then transferred to a −80°C freezer until processed for qPCR. One half of the biopsy samples were extracted using the Qiagen RNeasy 96 kit, after bead-beating in the Qiagen TissueLyser II using a 5mm Zirconium Oxide bead. RNA quality was measured on the Agilent BioAnalyzer 2100, cDNA was synthesized using the Invitrogen SuperScript IV First Strand Synthesis kit and 8 μL RNA was used for each reaction. The samples were reacted with the Taqman Fast Advance Master Mix. Real time PCR was performed with the following Taqman probes (ThermoFisher, Carlsbad, CA) GAPDH as housekeeping gene, TNF-α and NOS2 for inflammation and ARG1 and IL10 for remodeling (see probes in Table 2). Samples were analyzed by the 2^−ΔΔCT^ method.

**Table 2.** Taqman Gene Expression probes

| Gene | Taqman assay and fluor |
| --- | --- |
| GAPDH | Mm99999915_g1, VIC |
| NOS2 | Mm00440502_m1, FAM |
| TNF | Mm00443258_m1, FAM |
| IL10 | Mm01288386_m1, FAM |
| ARG1 | Mm00475988_m1, FAM |

### Mechanical testing

Dog-bone shaped tissue templates were measured for thickness, then placed in a unidirectional tensile testing device (UTTD) (Figure 1). The skin samples were marked with one dot above and one dot below the area wounded in surgery, which was centered in the gauge. The tissue was then placed into the UTTD in a water bath filled with fresh PBS at 37°C. The software program PASCO Capstone (Roseville, CA) was used to capture video and images every 0.5-1 second during the skin pull, and the force was collected simultaneously. Strain analysis measured the length between the two dots from each image captured until the tissue ruptured and was analyzed using NIH ImageJ, v1.52a (Bethesda, MD). Max stress was calculated using equation b:

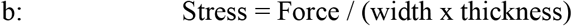

**Figure 1:**
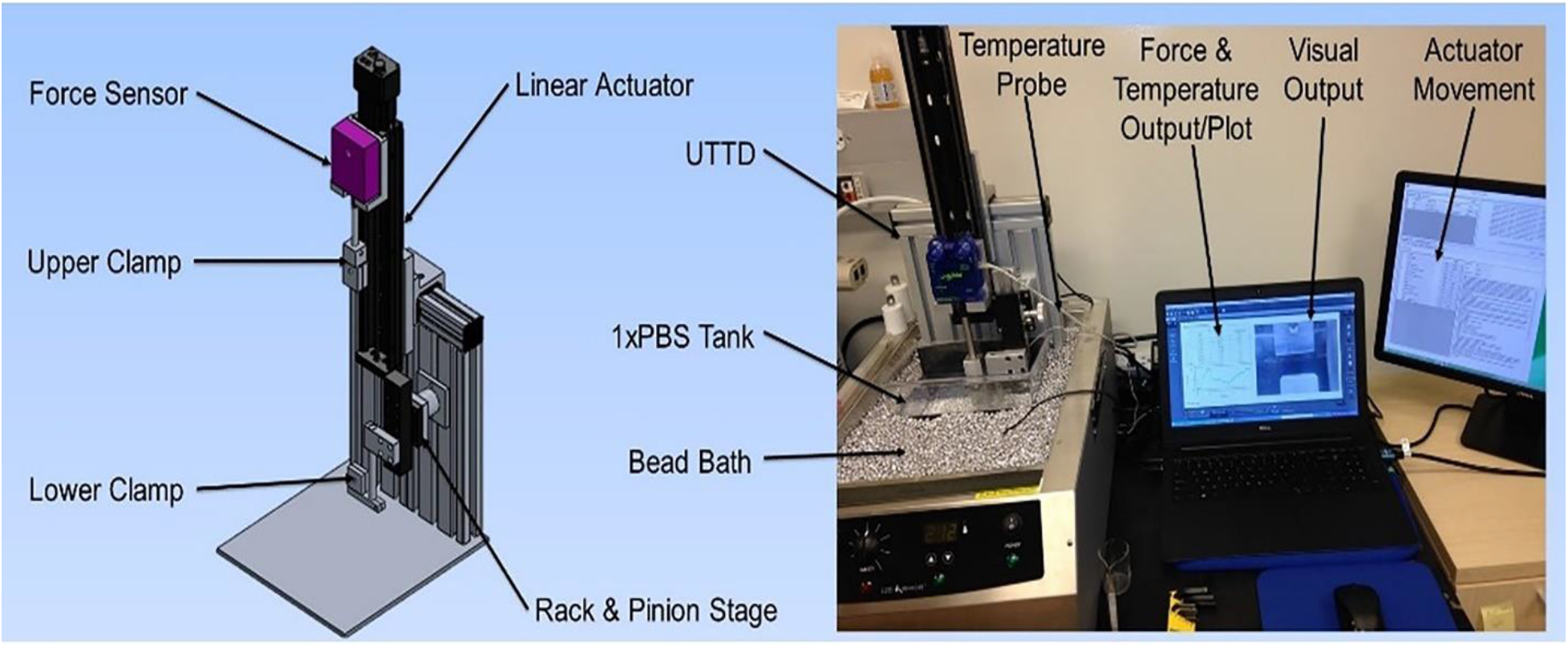
Left Uniaxial Testing Device (UTTD). Right, Assembled device and graphic user interface.

Max strain was calculated using equation c:

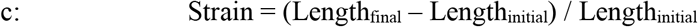

Elastic modulus was calculated using equation d:

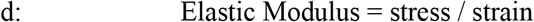

### Statistical analysis

Statistical analysis of wound closure rate, histological analysis, gene expression, and mechanical testing was performed using RStudio. The results of wound closure rate were analyzed using a Kruskal-Wallis test along with a Dunn’s post-hoc test. An ANOVA was used to analyze the mechanical property data and where the data did not meet the criteria for an ANOVA, a Kruskal-Wallis test was used. Histological analysis and qPCR were analyzed by using a multi-way ANOVA. Statistical differences were determined by p<0.05.

## Results

### Wound Closure

The data from the 14 and 28-day studies were combined for wound closure analysis. Percent wound closure for Control and WHD were significantly greater than Oasis from day 6 to day 16 (p < 0.05) at 6-13% over Oasis (Figure 2). After 20 days there was no difference among the three treatment groups. From day 14 through day 28 the average percent closure of WHD was 2-5% greater than control, though not statistically significant.

**Figure 2:**
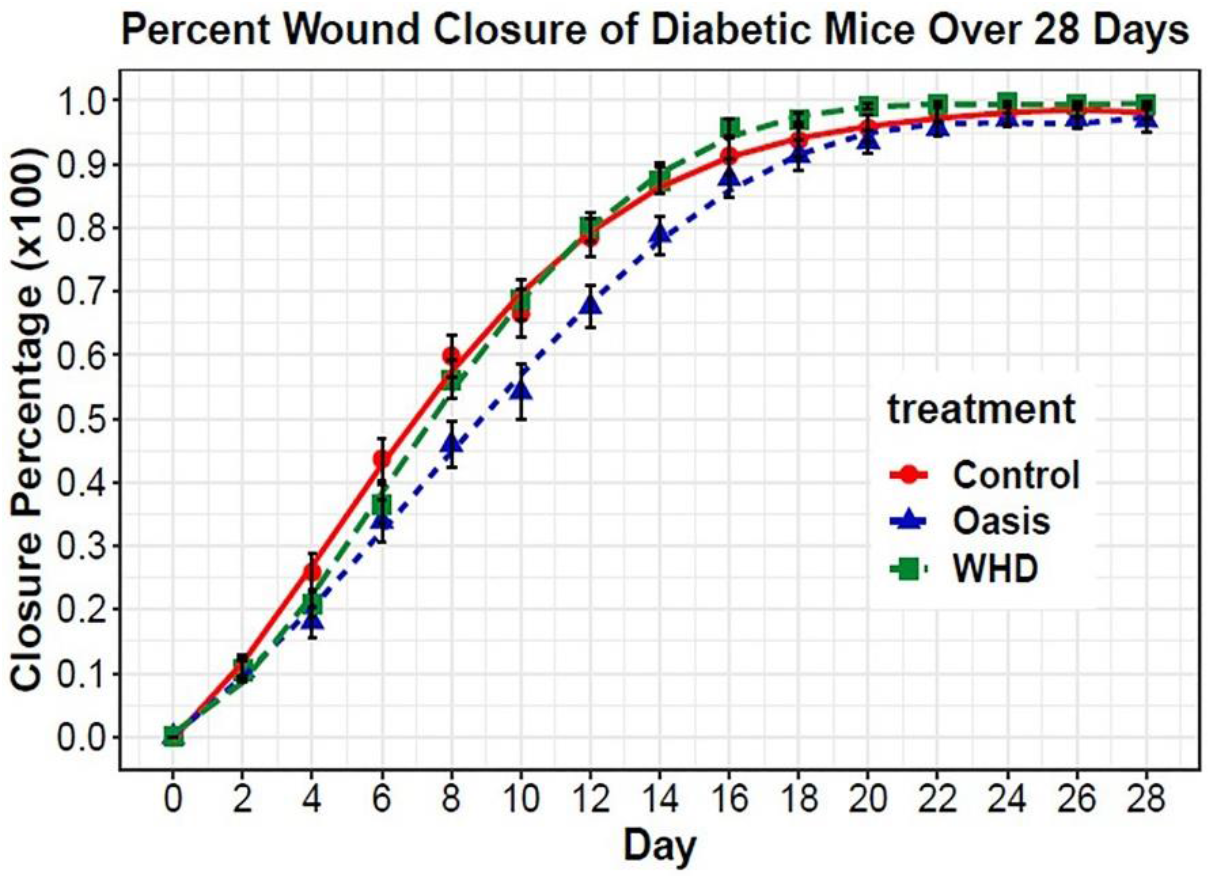
Polynomial regression analysis among treatment groups over 28 days. Percent wound closure was standardized to day 0 for each succeeding measurement. Shown values are means ± SEM.

No differences in sex were present as determined by multi-way ANOVA analysis (p = 0.08), therefore sex was removed as a variable. The wound closure data provides evidence that WHD-treated wounds achieve faster wound closure (most notable in days 6-16 of healing), compared to Oasis treatment in diabetic mice (Figure 3).

**Figure 3:**
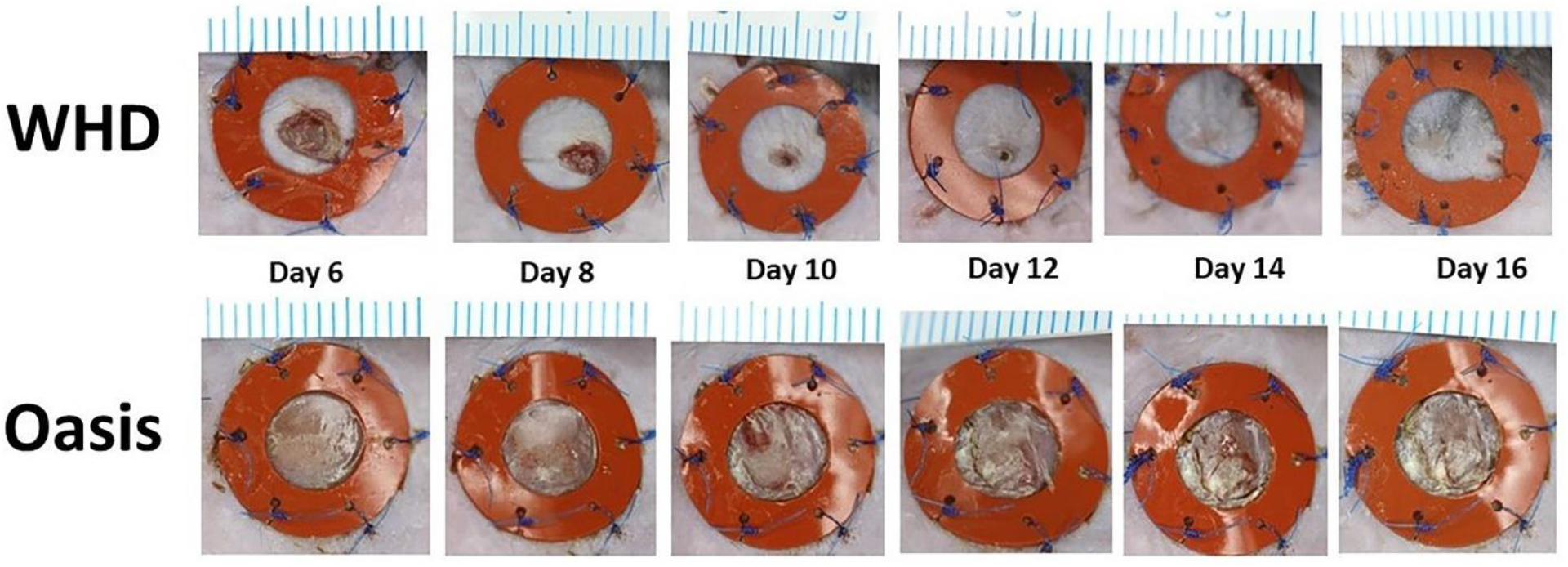
Gross images of full thickness splinted wounds in a diabetic murine model. Days 6-16 of healing demonstrate accelerated wound closure in the WHD treatment group compared to the Oasis Matrix Scale: each line = 1mm

### Histology and Immunohistochemistry

Histologically, wounds treated with the WHD demonstrated a greater degree of healing compared to the Oasis Wound Matrix or control treatment groups. The remodeled skin organ (epidermis and dermis) in WHD-treated animals exhibited well-formed new dermis, epidermis and accessory structures in the organ such as hair follicles and more closely resembled the native unwounded skin in the animal compared to Oasis-treated animals (Figure 4).

**Figure 4:**
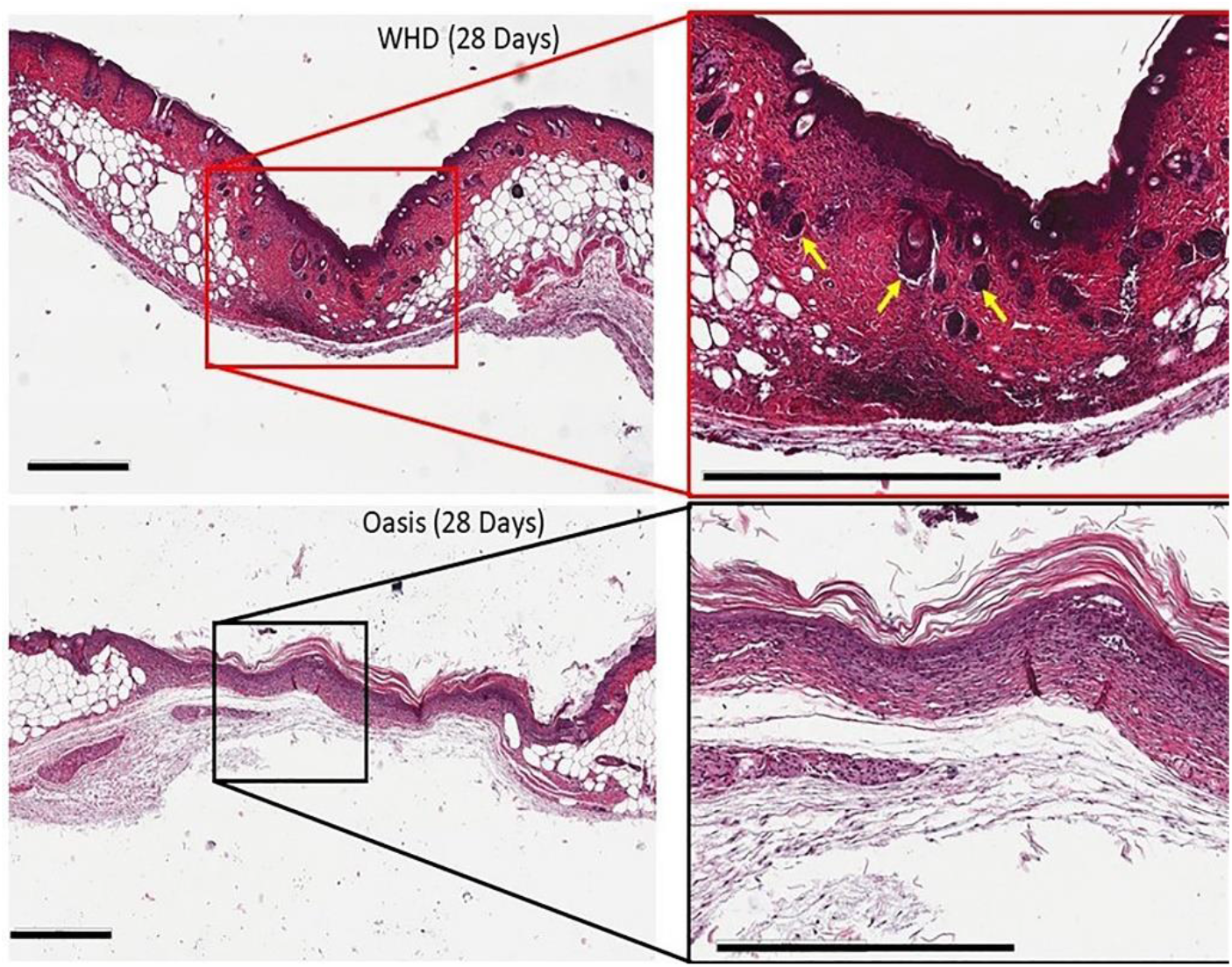
Hematoxylin and Eosin stained samples from WHD and Oasis Wound Matrix-treated wound sites after 28 days of remodeling. WHD-treated wound sites demonstrated wound regeneration where the skin organ and tissue had evidence of functional restoration with the presence of hair follicles (yellow arrows). Scale bars = 600 μm.

There was large variability between animals and between and within treatment groups and sex when quantifying IHC targets. Analysis of elastin showed no differences between treatment or sex at 14 or 28 days; however, there was a notable and significant (p < 0.001) increase in total elastin in the wound bed from 14 to 28 days (Figure 5), suggesting an increased elastin deposition as wound healing progresses, irrespective of treatment or sex.

**Figure 5:**
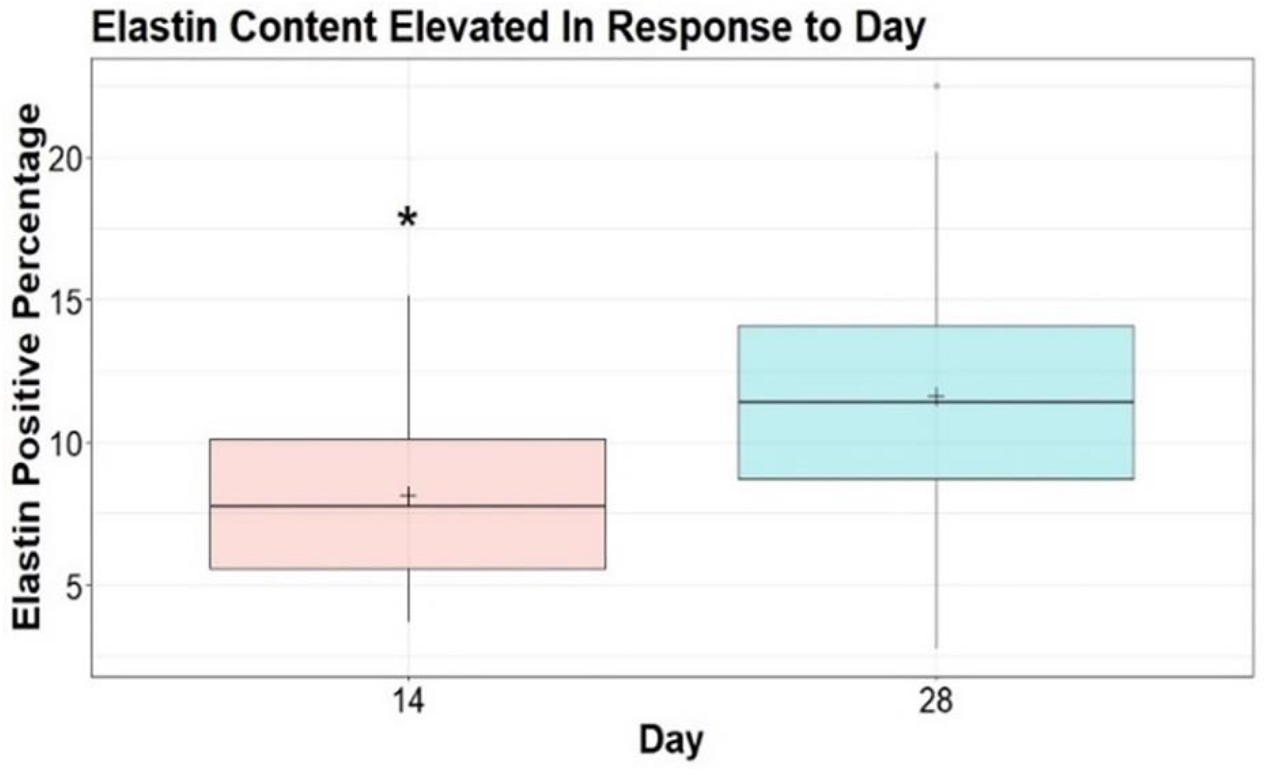
Analysis of elastin positive staining percentage between day 14 and 28. Asterisk represents significantly different group means where p = 0.0003. Comparison was conducted using a two-sided t-test with equal variance (Levenne’s Test; p = 0.08). Means are illustrated by crosses.

Microvessel analysis of the CD31 antigen, at 14 and 28-days post wounding, identified no differences between treatments, sex, or day (p = 0.94; p = 0.84; p = 0.45, respectively) (Figure 6). While there was no significant difference in the amount of microvasculature per area analyzed, there was less area to measure in the WHD sites at day 28 due to the progression of the healing. (Figure 7 right). Both treatments had a large amount of microvasculature present at day 14. (Figure 7 left).

**Figure 6:**
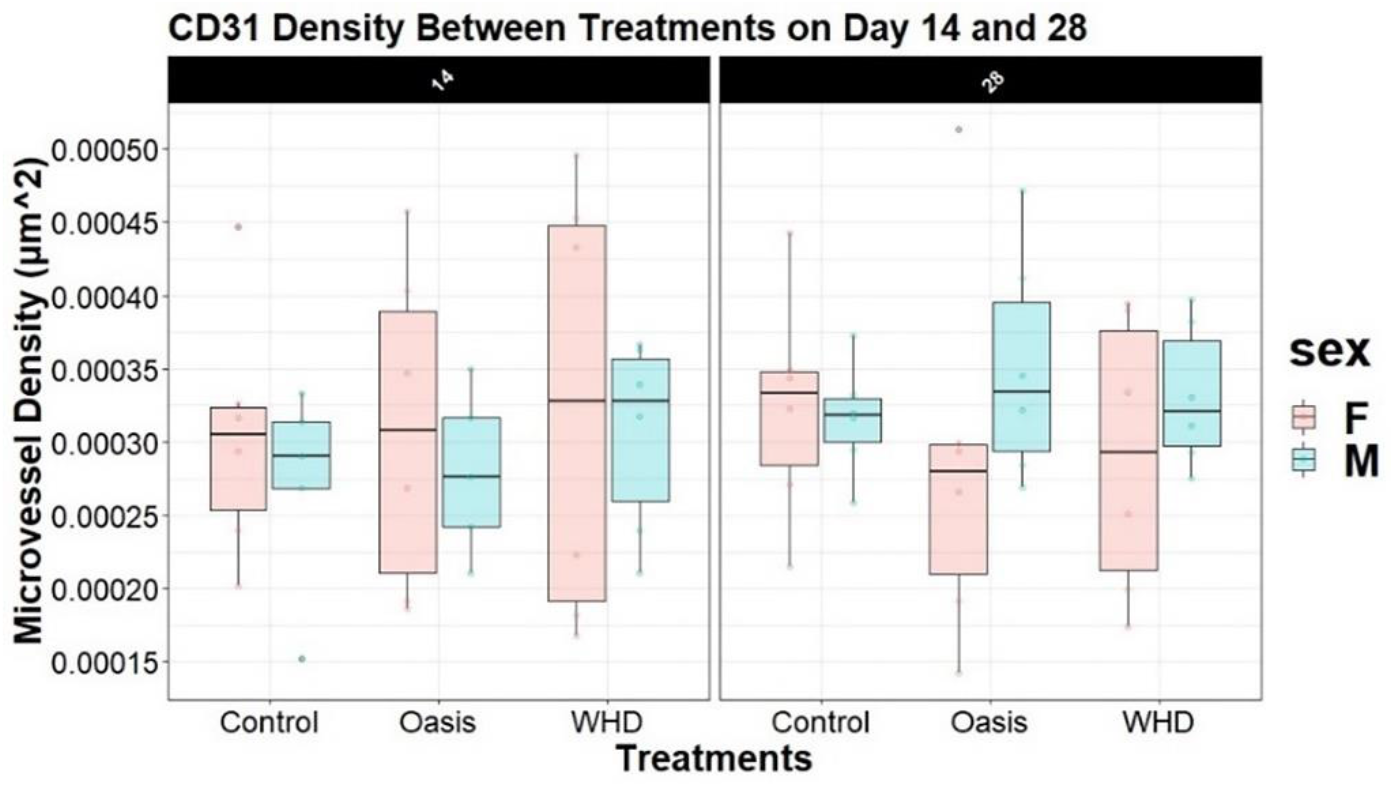
Microvessel density (CD31) faceted by day, treatment, and sex.

**Figure 7.**
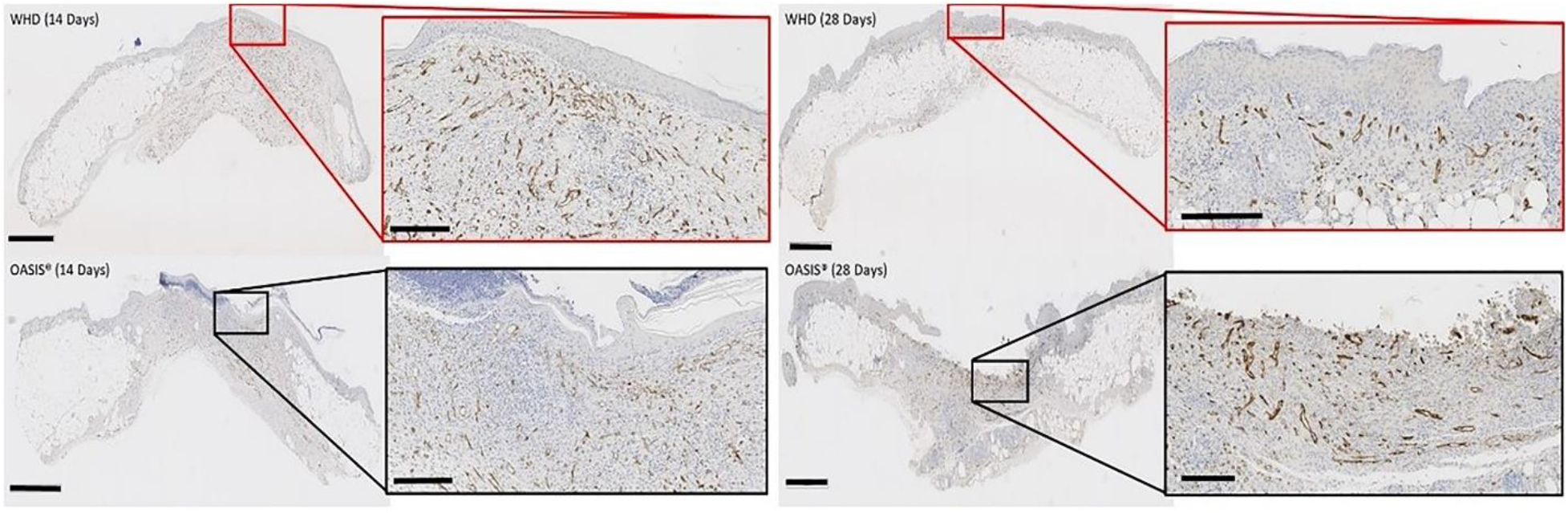
CD31 reacted tissues used to quantify the presence of microvasculature in the wounded tissue sites. Both treatments had a large amount of microvasculature present at 14 days (Left). While there was no significant difference in the amount of microvasculature measured per square micron, the area measured in the WHD was smaller due to the progression of wound healing in the 28-day samples (Right). Macro-image scale bars = 1mm, micro-image scale bars = 200 μm.

Inflammation was evaluated indirectly using CD163-reacted tissue samples. CD163 is an M2-like macrophage phenotype marker associated with the transition of the inflammatory stage and the proliferative and remodeling phase; therefore, comparisons between treatment and control groups were performed for the inflammatory comparisons (Hu et al., 2017). A multi-way ANOVA analysis of the variables sex, day, timepoint and treatment was performed iteratively to assess macrophage density in the wound beds identifying “treatment” as the sole variable impacting analysis. A one-way ANOVA on the final model suggested at least one treatment mean was different (p = 0.04, Figure 8). A Tukey pairwise t-test provides evidence that WHD treatment resulted in significantly less CD163^+^ macrophage presence when compared to Oasis; however, no differences were detected between control and WHD. At day 14 Oasis elicited a greater macrophage response in the wounded tissue compared to the WHD. By day 28 the macrophage response of the wounds treated with Oasis had decreased notably. The wounds treated with the WHD had a lower expression of CD163^+^ cells at both timepoints (Figure 9).

**Figure 8:**
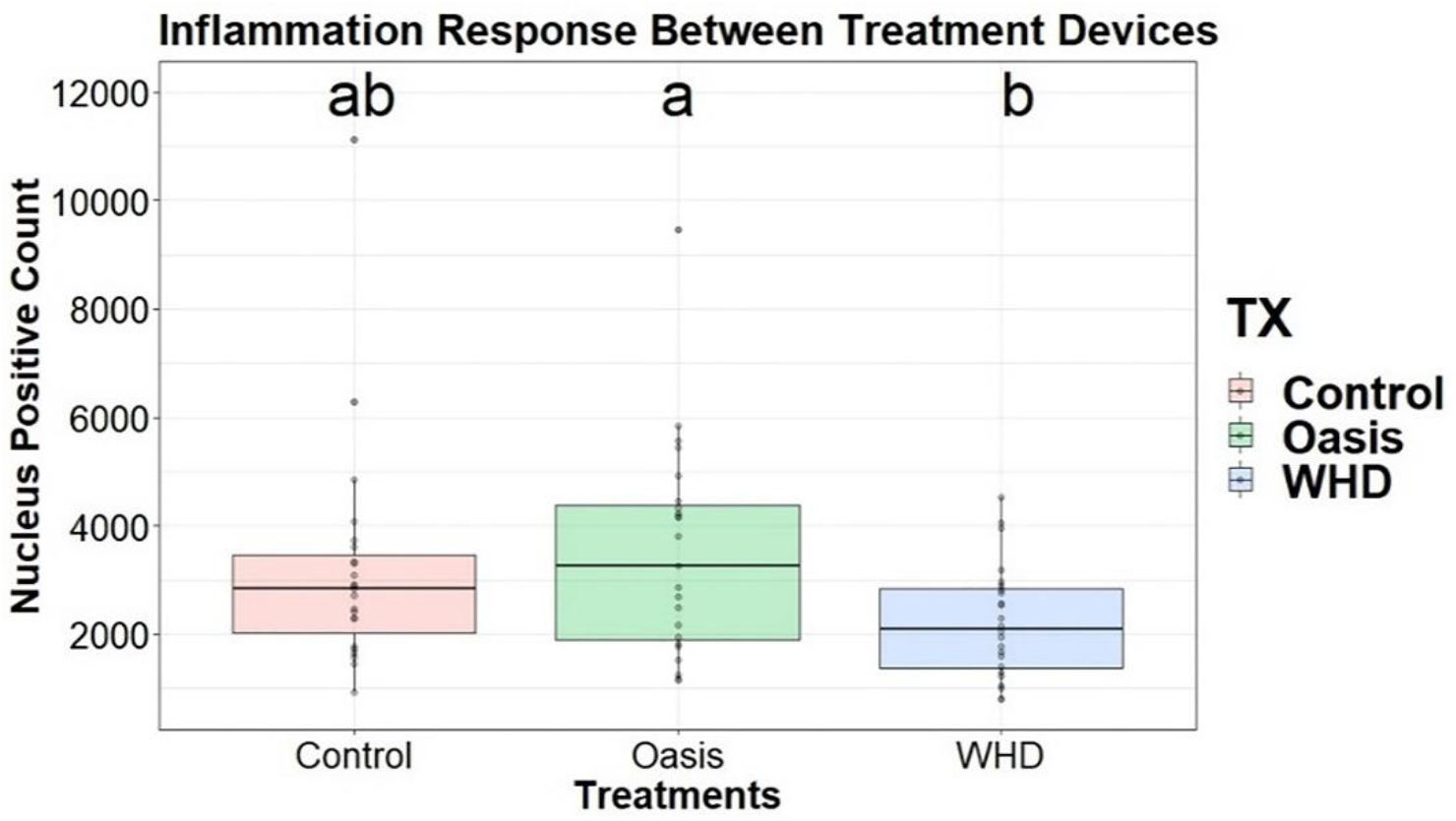
CD163 positive stain count compared across treatments. Compact letter display illustrates significant differences between treatments when p-value < 0.05. WHD treatment had significantly less macrophage presence during healing compared to Oasis.

**Figure 9.**
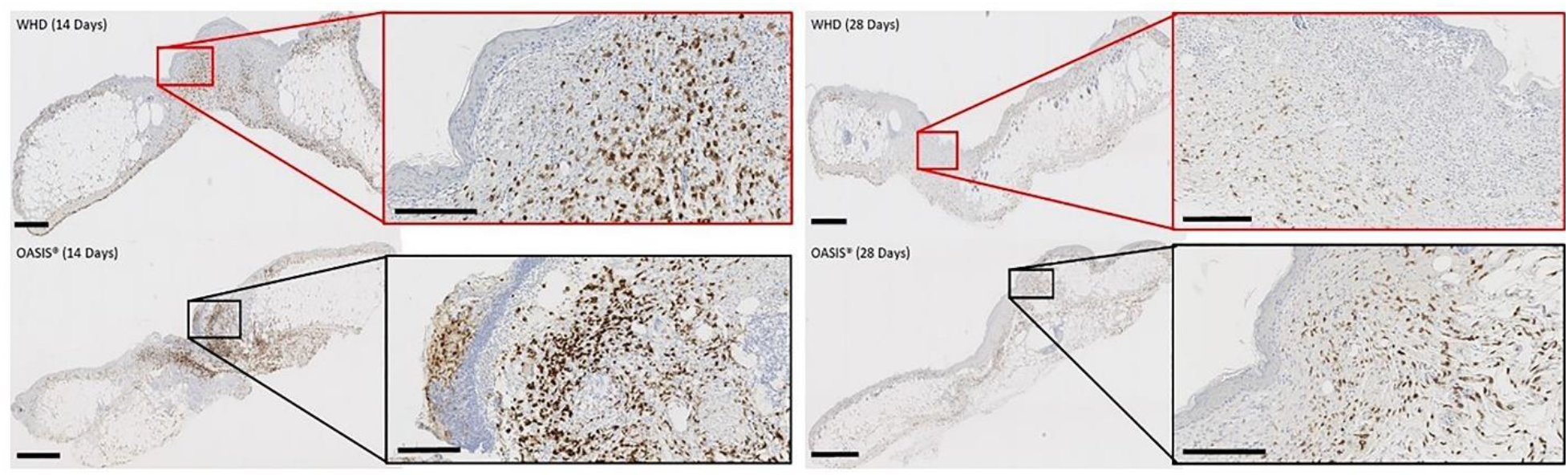
Phase I Immunohistochemistry (CD163) highlighting tissue macrophage presence in remodeled wounds at 14 days (left) and 28 days (right). The WHD demonstrated lower levels of tissue macrophage presence compared to Oasis.

### Gene Expression

The presence of genetic signals relating to specific phases of wound healing were evaluated: TNF-α and nitric oxide synthase (NOS2) for the inflammatory phase and IL-10 and arginase (ARG1) for the remodeling phase, normalized to housekeeping gene GAPDH.

The data showed a wide range of expression across all variables. A multi-way ANOVA analysis provided sufficient data to conclude that there was no significant difference found between “sex” or “treatment” in target gene expression (p = 0.13; p = 0.61). However, at 14-days, there was a slight trend toward an increase in expression of ARG1 and NOS2 in the Oasis group for both sexes compared to control and WHD, which were relatively equivalent. There were no trends observed in the 28-day group, likely related to the variability seen among animals (Figure 10). Furthermore, expression levels of all genes were similar at days 14 and 28. These findings indicate that the treatment groups did not induce any wound healing-related gene expression changes for the assessed gene targets in the current study.

**Figure 10:**
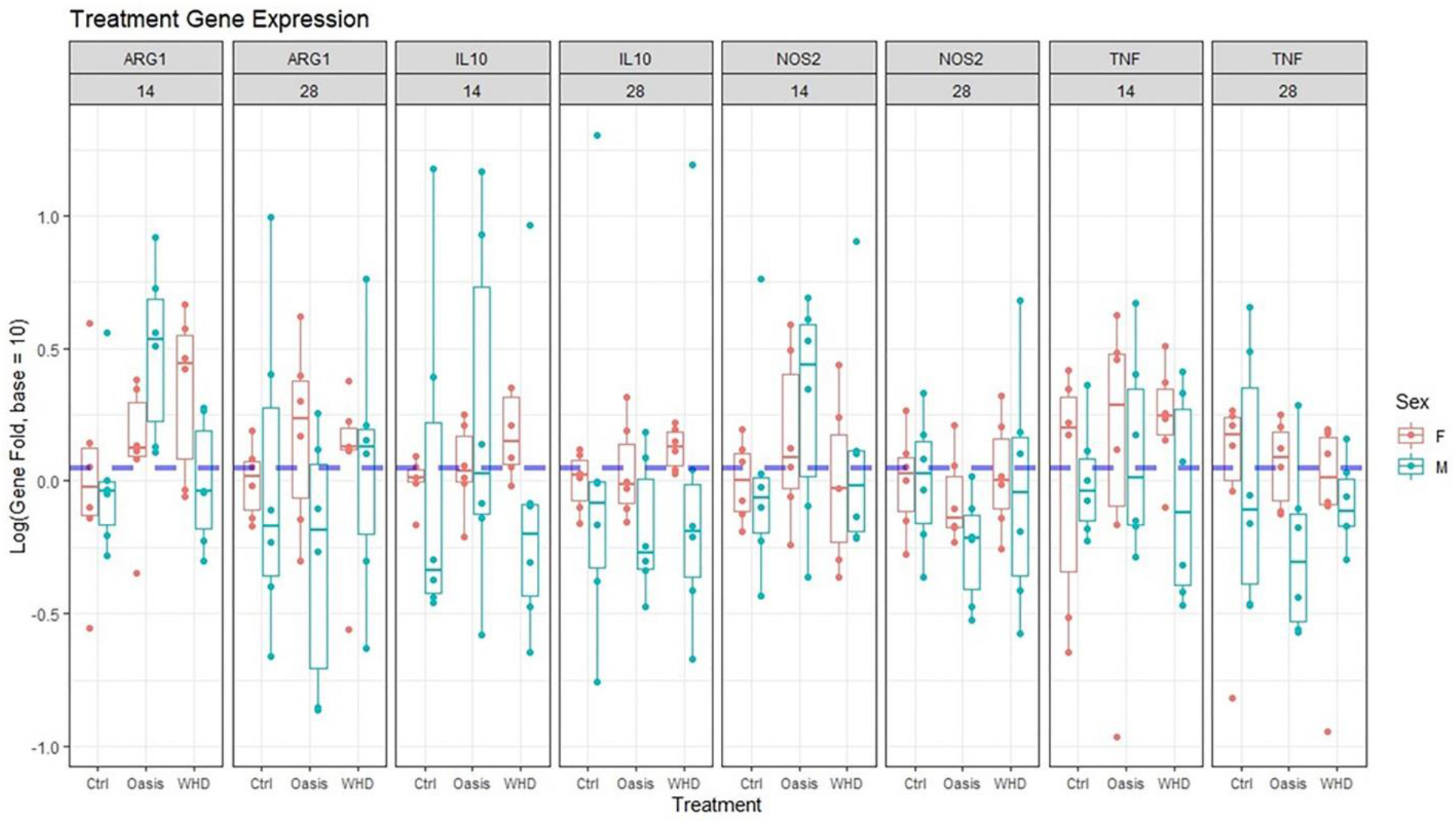
Treatment differences in gene expression faceted by sex, treatment, day, and gene. No significant differences were noted.

### Mechanical Testing

Mechanical testing was performed on remodeled tissue from WHD n=11, Oasis n=9, and Control n=12 treated wound sites. At Day 14, the Elastic Modulus for Oasis-treated wound sites was significantly lower than WHD and Control. From Day 14 to Day 28, there was a significant increase in Max Strain within all treatments groups (Table 3). At 28 days the WHD treated tissue demonstrated mechanics more similar to unwounded diabetic skin (Figure 11). Collectively, these data suggest that Oasis treatment produced a stiffer, less compliant remodeled wound tissue compared to control and WHD treated sites. Figure 11 demonstrates that the WHD withstood the greatest force prior to yielding with more than 0.4 MPa of stress compared to control before yielding; however, these data were not statistically significant due to the increasing variation as strain increased.

**Figure 11:**
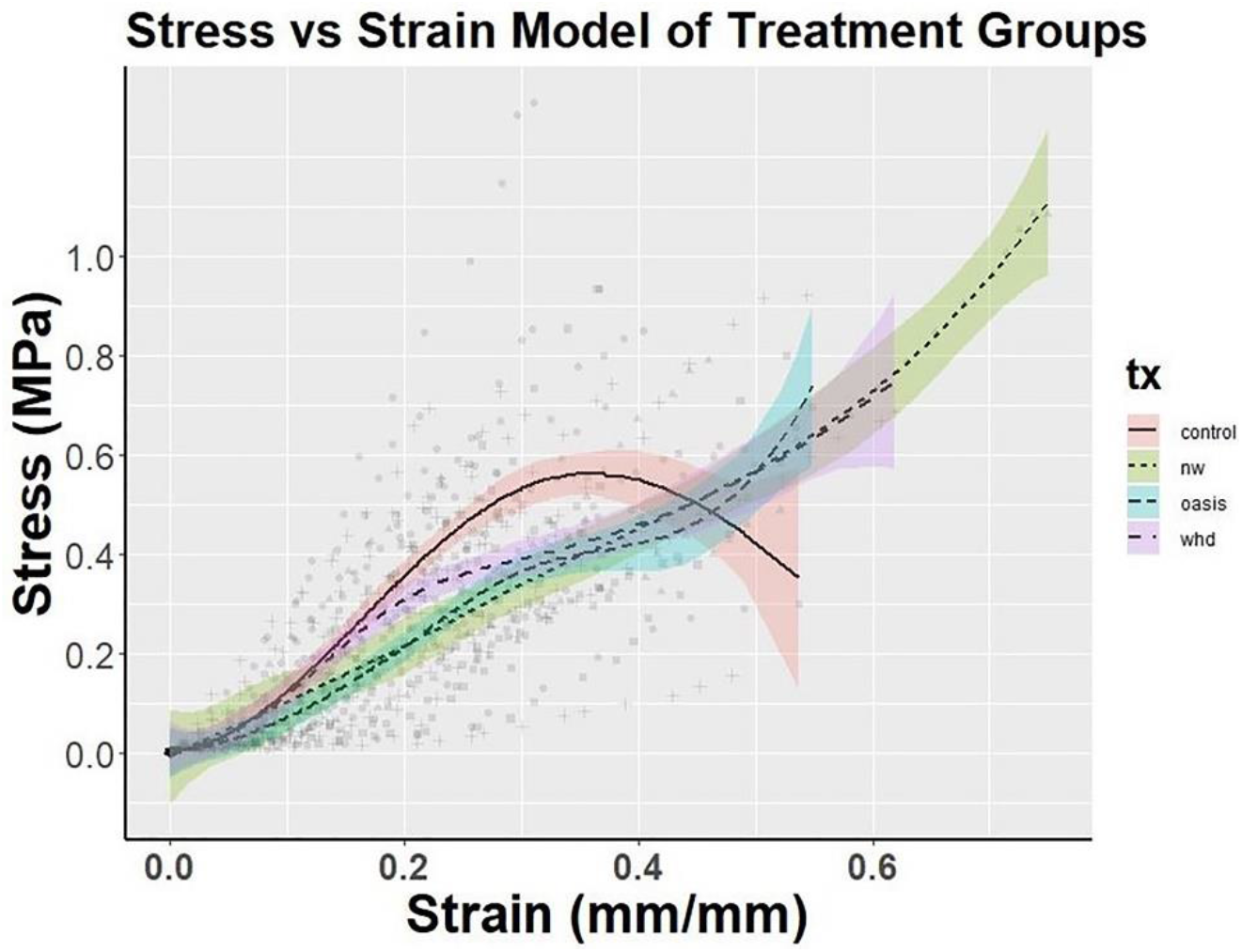
Stress vs. Strain analysis of excised murine skin for each treatment. Lines represent spline regression models with 95% confidence intervals. Data was divided into 3 intervals x < 0.2, 0.2 ≤ x ≤ 0.3, and x > 0.3 applying a cubic prediction model to each section. Although not statistically significant, due to the amount of variation introduced as strain increases, WHD withstood more than 0.4 MPa of stress before yielding compared to control. NW group = non-wounded skin.

**Table 3:**
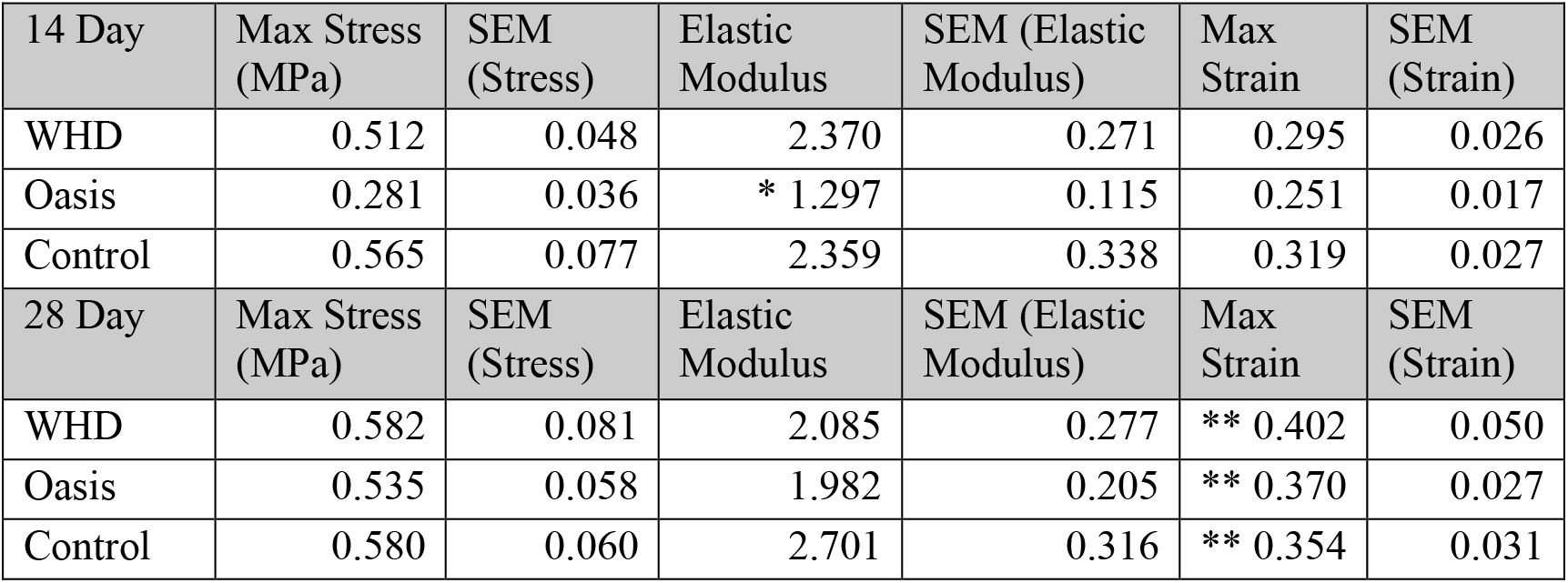
Mechanical testing results of Max Stress, Elastic Modulus, and Max Strain from Day 14 and Day 28 mice. Sample sizes for each treatment were: WHD n=11, Oasis n=9, Control n=12. Elastic Modulus in Oasis-treated wounds was significantly lower than WHD and Control at Day 14 (*, p<0.05). Max Strain significantly increased in all treatments from Day 14 to Day 28 (**, p<0.05).

| 14 Day | Max Stress (MPa) | SEM (Stress) | Elastic Modulus | SEM (Elastic Modulus) | Max Strain | SEM (Strain) |
| --- | --- | --- | --- | --- | --- | --- |
| WHD | 0.512 | 0.048 | 2.370 | 0.271 | 0.295 | 0.026 |
| Oasis | 0.281 | 0.036 | * 1.297 | 0.115 | 0.251 | 0.017 |
| Control | 0.565 | 0.077 | 2.359 | 0.338 | 0.319 | 0.027 |
| 28 Day | Max Stress (MPa) | SEM (Stress) | Elastic Modulus | SEM (Elastic Modulus) | Max Strain | SEM (Strain) |
| WHD | 0.582 | 0.081 | 2.085 | 0.277 | ** 0.402 | 0.050 |
| Oasis | 0.535 | 0.058 | 1.982 | 0.205 | ** 0.370 | 0.027 |
| Control | 0.580 | 0.060 | 2.701 | 0.316 | ** 0.354 | 0.031 |

## Discussion

Chronic non-healing, or slow to heal wounds present a global health concern. A concerning subtype of these chronic wounds are diabetic foot ulcers (DFUs). In a 1999 US study, an estimated average outpatient cost of treating one DFU event over a two-year period was calculated to be USD $28,000 (Ramsey, 1999). The increasing prevalence of sedentary lifestyle choices leads to diseases such as diabetes and obesity, which also contribute to the rise in these staggering wound numbers (Frykberg 2015, Sen, 2019). Based on these growing trends, novel wound-care therapeutics are essential to help reduce hospitalization re-admittance rates, decrease wound healing time, and prevent life altering amputations.

Our group has previously developed bioengineered scaffolds using human collagen and recombinant dermal tropoelastin in an architectural orientation that mimics the extracellular matrix. Therapeutics such as platelet-rich plasma, human adipose-derived stem cells, and growth factors that promote accelerated wound healing, have been previously incorporated into a WHD and tested in an *in vivo* mouse full thickness wound model. In these prior studies, expedited wound closure occurred compared to control groups (Machula. 2014 & Tabor, 2016). In the current research, we have shown that these electrospun WHDs alone (without stem cells or added growth factors) promoted faster cutaneous wound closure *in vivo* compared to commercial treatments, specifically the Oasis Wound Matrix. This current research focused on collagen’s contribution to the tensile strength of skin (Bailey, 1976), as well as elastin’s contribution to skin elasticity (Almine 2012). Furthermore, in adult wound healing, elastin is typically produced in small percentages, or not at all, compared to its production in juveniles scars (Almine 2012). Therefore, in the current study, an electrospun scaffold containing both type I collagen and tropoelastin, a dermal elastin precursor, was evaluated in a full thickness, splinted, diabetic, wound model, to assess wound healing outcomes with both collagen and elastin presence in the delivered wound healing scaffold, WHD. Since diabetic wounds are commonly associated with subsequent chronic infections, it is crucial for the wound to heal efficiently and obtain a proper mechanical barrier to moisture and microbial agents that can further delay the wound healing process due to improper scar formation.

Wound healing is a complex, series of molecular and cellular processes that work in concert to lead to resolution (regeneration) or repair (healing) (Clark, 1996). In wound resolution the tissue experiences regeneration of the extracellular matrix, cell population, and function of the tissue or organ. In contrast, wound repair is a healing response where scarring or fibrosis is present due to the lack of ability to fully regenerate the tissue. In the current study wound resolution and repair were characterized by evaluating closure rates of wounds and physiological characteristics of the injured tissue in a diabetic murine model. This study involved a standardized wound treatment model (control) and two treatment groups: an electrospun WHD, dual-protein (type I collagen and tropoelastin) and a commercially available porcine small intestinal submucosa (SIS) matrix, Oasis Wound Matrix. The safety and efficacy of the WHD was demonstrated through an IACUC approved, pre-clinical, full-thickness surgical study. In this study, percent closure rates were calculated; histopathology was assessed, gene expression changes were quantified by qPCR, and mechanical stress-strain testing was quantified.

The percent closure outcomes for control and WHD were significantly greater than Oasis from day 6 to day 16 (p < 0.05) at 6-13% over Oasis. From days 14 through 28, there was a statistically insignificant, though notable, increase in the average percent closure of 2-5% in the WHD treatment compared to control. Study results demonstrate that an electrospun WHD composed of human collagen I and tropoelastin applied to full thickness wounds, in a type II uncontrolled diabetic mouse model, resulted in accelerated wound closure compared to the commercial device, Oasis.

Hematoxylin and Eosin stained histological sections of control, WHD and Oasis-treated tissue demonstrated that the WHD-treated wounds had more complete organ regeneration (new skin) and closer epidermal/dermal resemblance to native murine epidermis. Follicular neogenesis was also visible in the WHD-treated wound sites based on the histological appearance of hair follicles within the wound margins when compared to the Oasis and control groups. These findings support the hypothesis that WHD-treated wounds advanced through the wound healing cascade at an accelerated rate resulting in wound resolution or skin regeneration, in comparison to the control and Oasis treated wounds. For example, the Oasis and control wounds had a fairly primitive wound covering, absent of well-organized and remodeled epidermis and dermis, with higher tissue macrophage presence and a lack of new hair follicles.

Immunohistochemistry was performed to evaluate angiogenesis, assessed with CD31, and inflammation, indirectly by the presence of tissue macrophages, characterized using CD163, in the newly remodeled wound sites. Although not statistically significant, at day 14, of the 28-day study, both Oasis and the WHD had a large presence of microvasculature in their wound margins, indicating angiogenesis had occurred within the newly remodeled wound sites. At day 28 the WHD treated wound margins were smaller in area due to the progression of the healing. Assessment of M2 macrophage presence demonstrated that the WHD treatment resulted in significantly lower tissue macrophage presence compared to Oasis. This finding suggests that Oasis-treated wounds may have still been in an inflammatory stage of wound healing where WHD-treated wounds had progressed to later stages of the wound healing cascade with less inflammation and more collagen deposition and wound resolution.

Genetic marker presence for specific phases of wound healing were also evaluated; however, no significant differences were seen in wound-healing related gene expression targets evaluated in the current study. This could be due to inherent variability between the animals, in which further testing, and a larger sample size may be required, or the specific markers (genes) of key phases of wound healing that were selected in the current study were not impacted by the different treatment modalities applied to the wounds.

An important goal for any wound recovery device is to rapidly close the wound and return the wounded skin to its native mechanical state. In the current study, the WHD in a splinted, full-thickness, diabetic murine wound healing model demonstrated an enhanced rate of wound closure, decreased tissue inflammation, skin organ regeneration, and a stronger and more durable remodeled tissue that more closely mimics native unwounded skin compared to the control groups. Additionally, the Stress-Strain analysis curve (Figure 11), illustrates that WHD-treated wounds, post-healing, demonstrated tissue mechanics that more closely mimic the non-wounded diabetic skin in this model. Collectively, these findings suggest that the WHD may aid in accelerating wound closure compared to the Oasis Wound Matrix and result in remodeled skin that more closely resembles the non-wounded native skin in mechanics and architectural structure.

## Acknowledgments

The authorship group would like to offer the following acknowledgment to The University of Arizona Genetics Core, University of Arizona, Tucson, AZ http://uagc.arizona.edu We appreciate your work and involvement in this project.

## Author Contributions

All persons who are authors certify that they have participated sufficiently in the work to take public responsibility for the content, including participation in the concept, design, analysis, writing, or revision of this manuscript.

## Author Disclosures

Robert S. Kellar PhD consults for Axolotl Biologix and is a scientific advisor for Protein Genomics.

Robert B. Diller PhD works for a regenerative medicine company, selling membrane products in the wound market.

Aaron J. Tabor PhD works for Axolotl Biologix.

Dominic D. Dominguez works for Axolotl Biologix.

Robert G. Audet works for Axolotl Biologix.

Tatum A. Bardsley works for Axolotl Biologix.

Alyssa J. Talbert has no disclosure to report.

Nathan Cruz has no disclosure to report.

Alison Ingraldi works for Axolotl Biologix.

Burt D. Ensley PhD is the owner of Protein Genomics.

